# Nanoscopic Origin of Enhanced Tyrosine-Mediated Water Ordering over Canonical Threonine Motifs in a Bacterial Ice-Nucleating Protein

**DOI:** 10.64898/2026.10.15.759343

**Authors:** Sankalp Jadhav, Ananya Debnath

**Author notes:** Electronic mail. Also at Department of Chemistry, Indian institute of Technology, Jodhpur,Karwad, Jodhpur, India-342037 (Sankalp Jadhav).

## Abstract

Bacterial ice-nucleating proteins (INPs) are known as the most efficient biological heterogeneous ice nucleators. Ice-nucleation ability is conventionally attributed to the threonine-based conserved motifs of *β* -helical INPs, whereas the role of other conserved motifs remains overlooked. To identify the motif with the highest hydration-ordering ability, a 16-loop model INP and its two mutants are simulated using all-atom molecular dynamics at 260 K, just below the ice nucleation temperature, using the CHARMM36m force field and TIP4P/Ice water model. Among four potential ice-nucleating sites present in four exposed surfaces of the *β* -helical INP, the glycine-tyrosine-glycine (GYG) motif-associated interface water (IW) exhibits the probability distribution of the Steinhardt bond-orientational and tetrahedral order parameters closest to that of the I_*h*_. The GYG arrange themselves into a ladder-like array of tyrosine hydroxyls, compatible with the I_*h*_-like lattice geometry, suggesting that this motif provides the IW a favorable template for ice-like water organization. The Y-to-phenylalanine mutation produces the largest disruption in IW order compared to the threonine mutations. Although the difference in entropy of threonine-associated IW and I_*h*_ is the least among all sites, ice-like hydration ordering near GYG may exceed prior assumptions for a threonine-based ice-nucleating surface. The ordering is collectively determined by lattice compatibility, entropy reduction, hydrophobicity, interfacial geometry, and sequence conservation. The study unfolds a molecular framework of biological ice nucleation and provides a foundation for the rational engineering of more efficient artificial ice-nucleators for applications in climate-related technologies.

## I. INTRODUCTION

The freezing of water to ice is of fundamental interest to many areas of science and technology, such as cryopreservation, food science, and the atmospheric sciences.^1–3^ Below 0° C, ice is the stable phase of water, yet due to kinetic barriers, pure water stays in the supercooled phase above the homogeneous nucleation temperature, which is as low as ∼ −38° C or ∼ 235 K.^4–6^ In contrast to this, ice nucleating protein (INP) derived from bacteria,^7–10^ can nucleate ice at much warmer temperatures −2° C (271 K) to −10°C (263 K), through heterogeneous ice nucleation.^10^ Thus, INP is known as one of the most efficient biological ice nucleators. INPs promote ice nucleation in clouds, specifically at low altitudes, due to their extreme environmental resilience, lower biological hazard, and abundance.^11,12^ Thus, INPs play a major role in the hydrological cycle and global climate and have immense applications in cryopreservation, agriculture, and food science. Despite years of extensive research and the importance of INPs on global climate, the molecular origin of ice nucleation by INPs remains elusive.

The central function of INPs is to template water molecules into an ordered, ice-like geometry, thereby overcoming the kinetic barrier that otherwise suppresses nucleation.^13–16^ INP-assisted nucleation leads to faster stabilization of the resulting polycrystalline ice composed of hexagonal ice I_*h*_ and cubic ice (I_*c*_) as compared to nucleation in pure water.^17^ Homogeneous nucleation produces multiple ice nuclei dispersed throughout the liquid, which subsequently grow and merge, resulting in an ice structure with a greater number of grains. In contrast, heterogeneous nucleation on the INP begins with the formation of a stable nucleus at the INP surface, which then grows continuously into the surrounding liquid, producing an ice structure with fewer grains. The faster stabilization of the heterogeneous ice structure is attributed to the reduced nucleation barrier associated with INP-mediated nucleation, thereby facilitating more rapid ice growth and grain-structure stabilization.^17^ The complete experimental structure of INP is not available due to the huge size (∼ 130 kDa) of the protein, its localization in the membrane, and the lack of experimentally resolved crystal structures.^18,19^ Early homology modeling suggests a central repeating domain (CRD) comprising ∼81% of the total sequence. The CRD has been proposed to contain the ice nucleation site of INPs, and it contains 16 amino acid tandem repeats, each following the well-established conserved sequence GYGSTXTAXXXSXLXA, with X denoting a non-conserved residue, forming an antiparallel *β* -sheet.^15^ The threonine-X-threonine (TXT) motif and serine-X-T (SXT) motif of each loop lie on the opposing sides of the flat surface of the *β* -helix of the INP. Using a fully rigid, united-atom representation of the INP, simulations of INP show that the two T residues of the TXT motif project outward from the strand at a spacing similar to the oxygen spacing of the basal and primary prismatic planes of the ice lattice.^20^ The matching lattice geometry is found to be the key mechanism for ice binding by TXT. SXT is proposed as another ice-binding site that nucleates ice with comparable efficiency to TXT, through an anchored-clathrate motif.^20^ However, the simulations do not consider the side-chain rotation, backbone fluctuations of the protein, and protein–water hydrogen bonding during the nucleation trajectories. In addition, nucleation is observed only after rescaling the INP loop geometry to match the lattice spacing of a different ice-binding protein (antifreeze proteins (AFPs)), as the native INP spacing does not produce ice nucleation under the simulation conditions used. While the rigid, coarse-grained approach is well suited to isolating the geometric lattice-matching mechanism, it leaves open the question of whether protein flexibility and the native (un-scaled) INP structure can alter the ice nucleation mechanism. This motivates the all-atom, flexible treatment adopted in the present work.

The ice nucleation efficiency of INPs over AFPs is attributed to a greater reduction in the rotational entropy of the TXT-associated IW than in supercooled water, as determined using all-atom molecular dynamics simulations.^21^ The results further suggest that the threonine ladder alone may not be sufficient for ice nucleation, and that a specific rigid structural context is likely needed to impart order to the surrounding water. Together, these studies^15,20–22^ essentially project the TXT motif as the principal candidate for the ice-nucleating surface of bacterial INPs, supported by its close lattice match to ice, its strong sequence conservation across INP homologs, and its ability to promote entropy reduction to the associated water.

Sum frequency generation (SFG) spectroscopy measurements demonstrate that increased IW ordering alone is insufficient to explain the high ice-nucleation activity of bacterial proteins; instead, their supramolecular interactions are critically required for this function.^23^ However, SFG does not provide information on ice-like orientational order, but captures mostly the orientational order of the bond vector with respect to the surface normal. SFG combined with molecular dynamics simulations and 2D infrared spectral analysis shows that INPs reorient at lower temperatures and attain a configuration where the ice-nucleating sites of the protein are more parallel to the water-air interface. This reorientation can enhance water interactions, generate a “flat, carpet-like geometry”, and thereby maximize available ice-nucleating sites for water.^19^ A recent experiment using high-speed cryo-microscopy provides evidence of freezing favored by the hydrophobic interface of the INPs.^24^

Similar to TXT or SXT, the GYG motif is well conserved across INP repeats,^22^ and has primarily been studied for its role in INP dimerization.^15,18^ Its potential contribution to ice nucleation itself has been overlooked so far, probably because, as a single planar tyrosine layer rather than a tandem TXT-type array, it is not initially expected to template ice in the same way. Although the role of tyrosine residues in INP has not been explored earlier, linearly stacked tyrosine residues have been identified as ice-binding motifs with efficiency between moderately active and hyperactive AFP.^25,26^ Therefore, a molecular-level understanding of the specific ice-nucleating sites on INPs, as well as the structural and interfacial factors governing their ice nucleation activity, remains a critical knowledge gap in the field. Motivated by this, the current study focuses on a systematic comparison of ice-like bond-orientational ordering, the tetrahedral order parameter, and the entropy of hydration layers across all four exposed faces of the INP monomer.

Hereafter, the sites are referred to as potential ice nucleating sites (PINS): glycine-tyrosine-glycine (GYG), threonine-glutamine-threonine (TQT), serine-glycine-serine-glutamate (SGSE), and serine-leucine-threonine (SLT). Remarkably, our results show that water near GYG is structurally more ordered compared to water near any of the other three faces, including the conventional ice nucleating site, TQT. The entropy calculated using the 2PT method shows that the entropy reduction of interface water (IW) with respect to supercooled water (SCW) is largest for TQT, followed by GYG and SLT. The mutations in the hydrophilic, water-facing residues of this INP PINS to hydrophobic ones reduce the structural order. The effect of mutations of INP PINS reveals that, in addition to the nature of the protein surface, geometry also plays a significant role. Our results establish that the reduction in entropy is not, by itself, sufficient to explain the emergence of IW ordering. Instead, we identify the GYG motif as a previously unrecognized molecular determinant of hydration structuring, capable of promoting an ordered interfacial water network to a greater extent than previously characterized threonine-based ice-nucleating motifs. These findings provide a molecular-level mechanistic framework linking specific protein sequence motifs to the structural organization and ice-nucleating propensity of interfacial water, thereby offering a rational basis for the computational design and engineering of enhanced artificial ice-nucleating proteins for climate-relevant applications.

## II. METHODS

### A. Molecular Dynamics Simulation Details

The well-investigated 16-residue sequence GYGSTQTSGSESSLTA^13,17,20^ of the InaZ INP of *Pseudomonas syringae* is utilized to generate the INP monomer in our simulation, where the key ice-nucleating residues are highlighted as the TXT motif. The optimized length of the protein for the highest ice nucleation efficiency of a single INP monomer is predicted to be 8 nm,^13^ which can be generated from 16 loop repeats at 247 K. Based on this, the all-atom INP monomer is constructed using AlphaFold-3^27^ by repeating the above mentioned sequence^20^ 16 times to form a 16-loop *β* -solenoid that comprises 256 residues in total (Figure 1a,c-g).

**FIG. 1.**
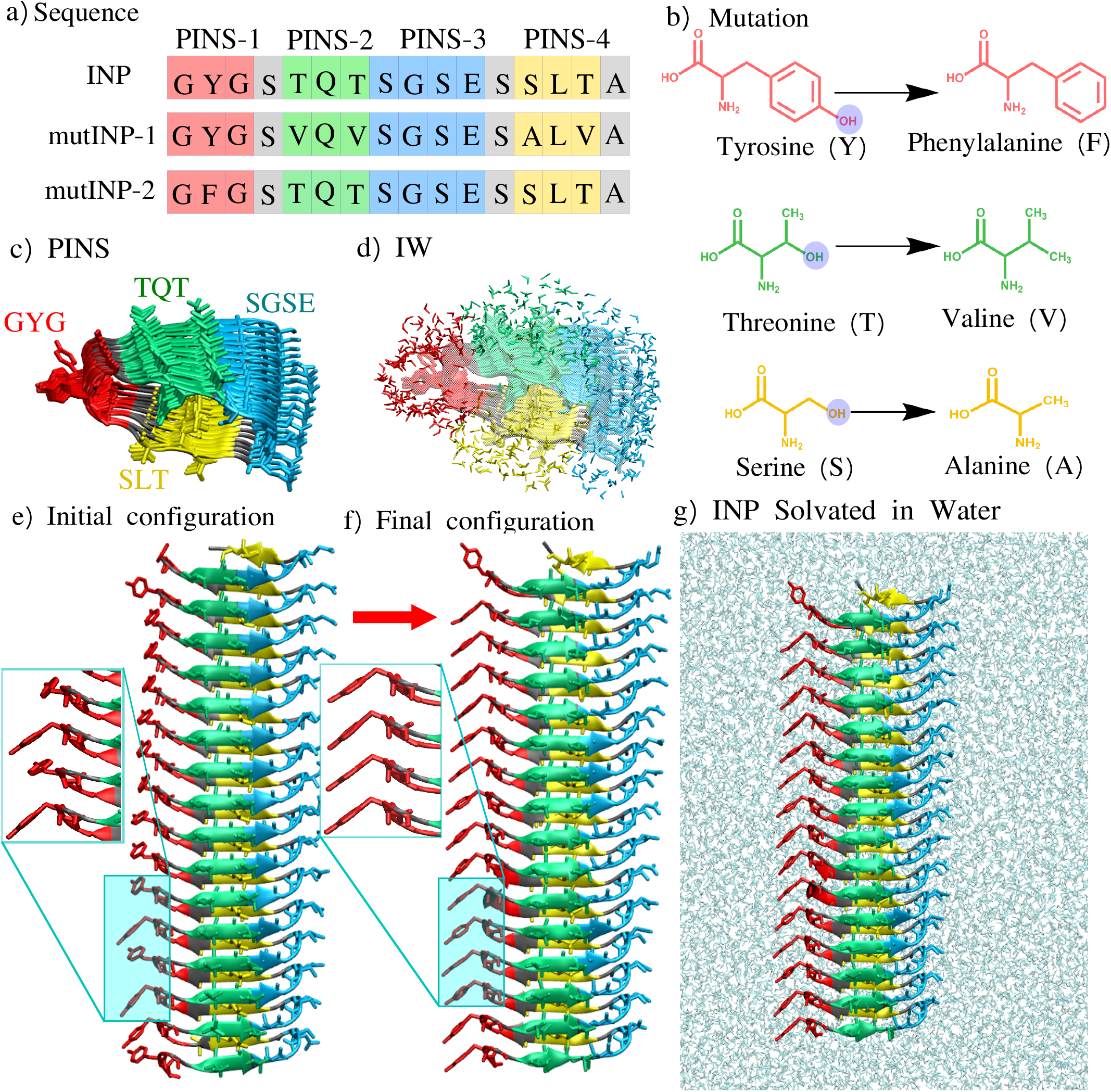
Sequence, structure, and mutation scheme of the ice-nucleating protein segment used in this study. (a) Sequence alignment of the wild-type INP repeat (containing the GYG, TQT, SGSE, and SLT motifs) compared with mutINP-1 and mutINP-2. (b) Residue-wise mutation tyrosine*→*phenylalanine, threonine*→*valine, and serine*→*alanine. (c) INP structure side view, colored by motif (GYG: red, TQT: green, SGSE: blue, SLT: yellow). (d) Water molecules in the first hydration shell of INP, colored using the same color code as PINS. (e,f) Initial and final simulated configurations of the *β* -solenoid INP monomer, with special emphasis on the random tyrosine arrangement in the initial configuration, which are stacked in the final configuration. (g) Full simulation snapshot of the INP monomer solvated in explicit water.

This approach captures the most essential ice-nucleating CRD while avoiding the expensive computational simulation of the full-length InaZ protein, which contains 78 loop repeats in its CRD.^28^ Within the used CRD, four surfaces of the *β* -solenoid protein are chosen as potential ice-nucleating sites (PINS) as: glycine-tyrosine-glycine (GYG) as the PINS-1, threonine-glutamine-threonine (TQT) as the PINS-2, serine-glycine-serine-glutamate (SGSE) as the PINS-3, and serine-leucine-threonine (SLT) as the PINS-4 (Figure 1a). To confirm the ice nucleation ability of the residues of the PINS, two mutant constructs are generated in which T is mutated to valine (V)^20^ and S is mutated to alanine (A) on the PINS-2 and PINS-4, and referred to as mutINP-1. Y is mutated to phenylalanine (F) and referred to as mutINP-2 (Figure 1a-b). All mutations are performed using the CHARMM-GUI server.^29–31^ The INP, mutINP-1, and mutINP-2 are solvated in TIP4P/Ice water,^32^ and the CHARMM36m force field^33^ is used to simulate INP and its mutated versions (see Table I). All protein-water systems are energy-minimized using the steepest-descent algorithm^34^ to remove unfavorable contacts. Next, position restraints of 400 kJ mol^−1^ nm^−2^ and 40 kJ mol^−1^ nm^−2^ are applied to the protein backbone and the side chains, respectively, and simulated for 125 ps in an NVT ensemble and for 5 ns in an NPT ensemble. Next, 1 ns NPT run is performed with the reduced position-restraint of 100 kJ mol^−1^ nm^−2^ on the protein backbone, and 40 kJ mol^−1^ nm^−2^ on the sidechains. In the next step, a 45 ns NPT simulation is performed without position restraints at 260 K using the Berendsen thermostat with a coupling constant of 0.5 ps where the pressure is maintained at 1 bar using a Berendsen barostat^35^ with coupling constant of 1 ps. Next, the NPT production run of 5 ns is carried out using the Parrinello-Rahman barostat^36^ with a coupling constant of 5 ps, and a velocity-rescale thermostat^37^ with a coupling constant of 0.5 ps. Previous studies show that the ice nucleation temperature without larger aggregation is around 263 K.^10^ Hence, 260 K is chosen for our simulation, which is below the ice nucleation temperature. The long-range electrostatic interactions are calculated using the particle mesh Ewald summation method^38^ with a cutoff of 1.2 nm and a grid spacing of 0.16 nm. van der Waals interactions are calculated using a cutoff of 1.2 nm. The simulation timestep for all cases is 2 fs. The trajectory is saved every 1 ps. For 2PT analysis, a 200 ps production run is performed using a timestep of 1 fs, and the trajectory is saved every 4 fs. The details of all systems have been mentioned in Table I.

**Table I.** Details of all systems simulated with equilibration and production run lengths.

| System | Water molecules | Box size (nm) | Equilibration (ns) | Production (ns) |
| --- | --- | --- | --- | --- |
| INP-monomer | 86228 | 13.8×13.8×13.8 | 51 | 5.2 |
| mutINP-1 | 86228 | 13.8×13.8×13.8 | 51 | 5.2 |
| mutINP-2 | 85070 | 13.8×13.8×13.8 | 51 | 5.2 |
| SCW | 3404 | 4.3×4.3×5.5 | 5 | 5.2 |
| $I_h$ | 2000 | 3.9×3.6×4.5 | 5 | 5.2 |

The supercooled water (SCW) is simulated with a total of 3404 TIP4P/Ice^32^ water molecules in a cubic box. An initial hexagonal ice (I_*h*_) configuration is created, consisting of 2000 water molecules, using the genice2 software.^39^ The system is energy-minimized using the steepest-descent algorithm.^34^ SCW and the bulk ice are equilibrated for 100 ps in the NVT ensemble, followed by 5 ns equilibration runs at 260 K using the Berendsen thermostat and barostat.^35^ Next, 5 ns-long NPT production runs are performed for each case using a V-rescale thermostat^37^ with a coupling constant of 0.5 ps and a Parrinello-Rahman barostat^36^ with a coupling constant of 5 ps. The system details and the run lengths are mentioned in Table I.

### B. Interface Water

To identify the structural and dynamical changes induced in water by INP, we have performed structural and dynamical calculations on water at the protein interface. For all the structural calculations (Steinhardt order parameter, tetrahedral order parameter), all the oxygen atoms of the water molecules within the first hydration shell of all the heavy atoms of side chains of PINS (0.32 nm for INP PINS and 0.45 nm for mutated INP PINS) are considered as interface water (Figure S1, Supplementary Material). For entropy calculations, all water molecules that continuously reside within 0.54 nm of heavy atoms of INP PINS are considered as interface water. The 0.54 nm cutoff corresponds to three hydration shells for INP PINS and two hydration shells for mutated INP PINS. The hydration shell cutoffs are chosen based on the radial distribution function (Figure S1, Supplementary Material).

### C. Steinhardt Bond Order Parameter

The local Steinhardt order parameter q_*l*_ for particle *i* is defined as^40^,

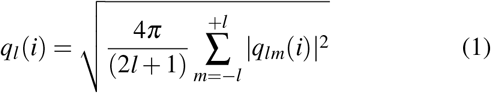

where the (2*l* + 1) components are computed as,

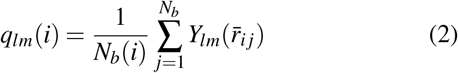

*l* is an integer parameter, *m* is an integer parameter that runs from *m* = −*l* to 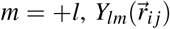 are the spherical harmonics calculated for the 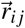 connecting the central particle *i* to the neighboring particle *j*, and *N*_*b*_(*i*) is the number of closest neighbors of the particle *i* within the given cutoff *r*. Because local crystalline structures are difficult to distinguish using Steinhardt order parameters, whose distributions smear out due to thermal fluctuations, the modified Steinhardt order parameter is calculated by averaging over the bond-order parameter of nearest-neighbor particles^41^ defined as

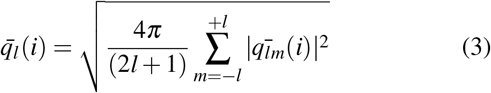

Where

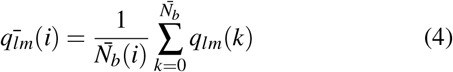

Here, the sum from *k* = 0 to 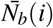 runs over all neighbors of particle *i* plus the particle *i* itself. Thus, to calculate 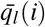 of particle *i*, one uses the local orientational order vectors q_*lm*_(*i*) averaged over particle *i* and its surroundings. While *q*_*l*_(*i*) holds the information of the structure of the first shell around particle *i*, its averaged version 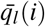 also takes into account the second hydration shell. The cutoff for finding the neighbor water oxygen atoms of the central atom *i* is set to *r* = 0.35 nm, which corresponds to the first hydration shell of water molecules. The only water molecules with *N*_*b*_(*i*) ≥ 3 are chosen for the calculation. Plumed-2.10.0 is used to calculate the Steinhardt bond order parameter^**?**^.

### D. Tetrahedral Order Parameter

The tetrahedral order parameter (q_*tet*_) is calculated as,^42,43^

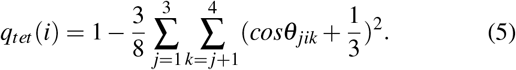

where *θ*_*jik*_ is the angle between central atom *i* and any two of the four nearest neighbors *j* and *k*. The q_*tet*_ is computed for all the IW near the PINS of INP and mutated variants, as defined in section II B. To calculate q_*tet*_, the neighbors of a central atom are defined as its four nearest atoms within a 0.35 nm cutoff. The four nearest atoms can be either water oxygen atoms or protein heavy atoms.

### E. 2PT Entropy

To understand the thermodynamics of water in hydration layers, we calculated the entropy of water near the PINS for INP, its mutants, and bulk water using the 2-Phase Thermodynamics (2PT) method.^44–48^ The 2PT method determines the thermodynamic properties of a system from its density of states (DOS) function. It decomposes the velocity and angular-velocity power spectra (density of states) into gas-like and solid-like components. The DoS *g*(*ν*) of a liquid is obtained by taking the Fourier transform of the mass-weighted velocity autocorrelation function (*C*(*t*)). The translational VACF is given as

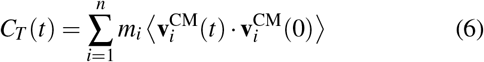

where *m*_*i*_ is the mass of the *i*^*th*^ molecule, 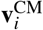 is the translational velocity of the center of mass of *i*^*th*^ water, and *n* is the number of molecules included in the summation. Similarly, the moment-of-inertia-weighted angular VACF is given by

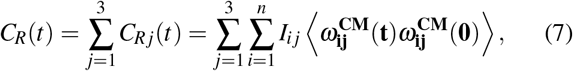

where *I*_*ij*_ and *ω*_**ij**_ are the *j*^*th*^ principle moment of inertia and angular velocity of water molecule *i*. The DoS *g*(*ν*) of a liquid is given as

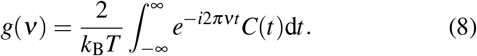

The DoS is partitioned into gas-like and solid-like components

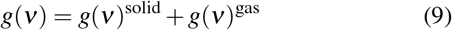

where the entropy of the system is calculated by weighting the individual density of states as

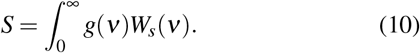

The solid-like DoS is treated using the harmonic oscillator (HO) model, while gas-like diffusion is described by the Enskog hard-sphere (HS) theory for translation and the rigid-rotor (RR) model for rotation. These contributions are then combined to obtain the total entropy of water.

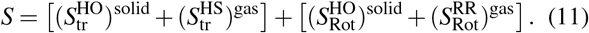

The model independently evaluates the translational and rotational entropy contributions. A detailed description of the method is mentioned in previous works.^45,48,49^ The 2PT method allows the translational and rotational entropy to be estimated directly from short MD trajectories without extensive sampling. A separate 200 ps production trajectory is generated with a 4 fs saving frequency, and the entropy is calculated for IW that continuously reside within 0.54 nm of the respective PINS for a 20 ps run length. The effective volume occupied by water molecules is calculated using Voronota-LT software.^50^ The reported entropy is averaged over five uncorrelated data sets.

## III. RESULTS AND DISCUSSIONS

### A. GYG Promotes Maximum Ice-Like Bond-Orientational Ordering of Interfacial Water among All PINS

The RDF shows long-range ordered structure and high solvent accessibility for GYG PINS compared to other INP PINS (Figure S1 of the Supplementary Material (SM)). For the mutated PINS, water molecules move away from the surface of PINS and do not show any long-range order. The probability density function (PDF) of the Steinhardt bond order parameters (SBOPs) calculated using equation (3) for interface water (IW) near the PINS of the INP monomer and its mutated variants is shown in Figure 2. The bulk water and I_*h*_ values of the 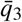 Steinhardt bond order parameter have been previously reported in the literature.^51^ For all cases of 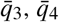,and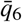, the location of the most probable peak of the PDF of the GYG is the rightmost among all PINS, closest to the most probable peak location of I_*h*_. This signifies that the GYG-associated IW exhibits the highest orientational order among the IW associated with all PINS, and surpasses the water ordering ability of conventionally known threonine-based motifs.^20^ The most probable peak locations of IW near TQT and SLT PINS are similar to each other, indicating that TQT and SLT sites have comparable ice nucleation abilities, in agreement with previous literature.^20^ SGSE residues exhibit the least order for all SBOPs. The overall trend from SBOPs for IW ordering of the INP monomer is GYG > TQT ≈ SLT > SGSE. The higher order for IW near GYG originates from the linearly ordered structure of the oxygen atoms present at the tip of the tyrosine (Y) residues that arrange like a ladder during equilibration of the monomeric INP from its initial configuration (Figure 1e-f). GYG imposing the highest structural order in the IW compared to conventional TQT or SLT motif is in agreement with the previous experimental prediction that the threonine-based motif does not induce greater ordering of water than other PINS.^52^ To confirm the effects of Y, T, and S in the proposed PINS on ice nucleation, a mutation study is performed at the GYG, TQT, and SLT sites. As previous studies do not indicate the role of SGSE in ice nucleation, and the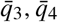, and 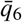 of the IW near SGSE have the least order among all PINS, SGSE is not considered for mutation. The 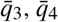,and 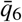of all mutated PINS shift away from the most probable location of the PDF of the I_*h*_ (Figure 3). The differences in peak location of PDF for the wild-type and mutated PINS are significant for 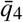 and 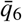 compared to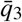, suggesting that 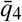 and 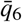 are better order parameters to understand the protein mutation effect on the structural order of the surrounding water. This confirms that the order of IW near GYG, TQT, and SLT originates from Y, T, and S. A significant reduction in the order is observed when GYG is mutated to GFG in mutINP-2 PINS (Figure 3a, d, g) due to the absence of a hydroxyl group in the F residues, indicating a major role of GYG in ice nucleation compared to the conventionally proposed threonine-based PINS.

**FIG. 2.**
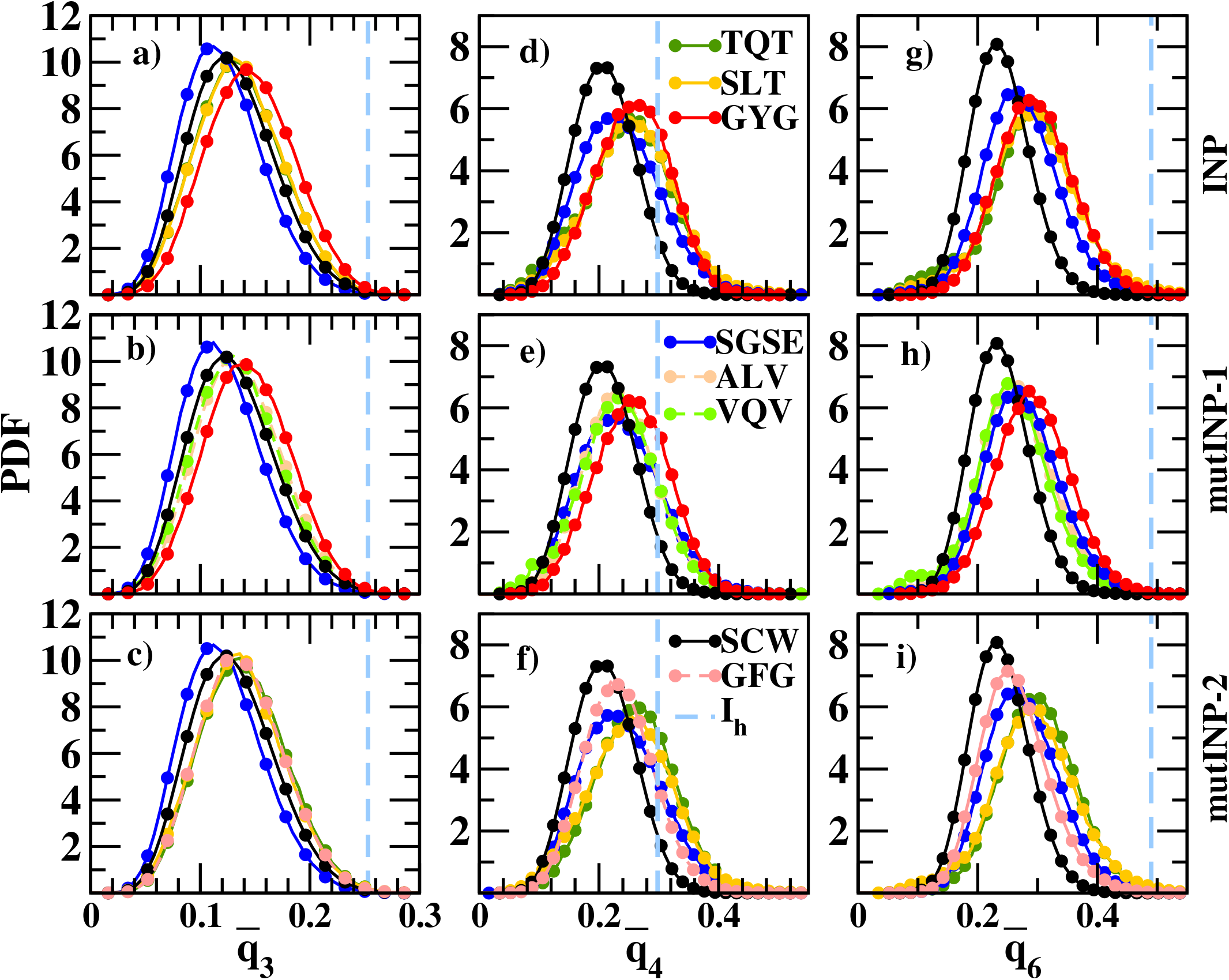
The probability density function (PDF) of 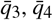, and 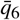 Steinhardt bond order parameter (SBOP) for the interface water (IW) near the PINS of the monomer. The most probable location of the PDF for hexagonal ice (I_*h*_) is shown with the dashed light blue line. The first (a, b, c), second (d, e, f), and third (g, h, i) columns represent 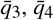, and 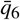 SBOPs, respectively. The most probable location for GYG shifts towards I_*h*_ more than for other PINS. The peak shifts towards that of the SCW once mutated to GFG, suggesting that GYG is the ice-nucleating site.

**FIG. 3.**
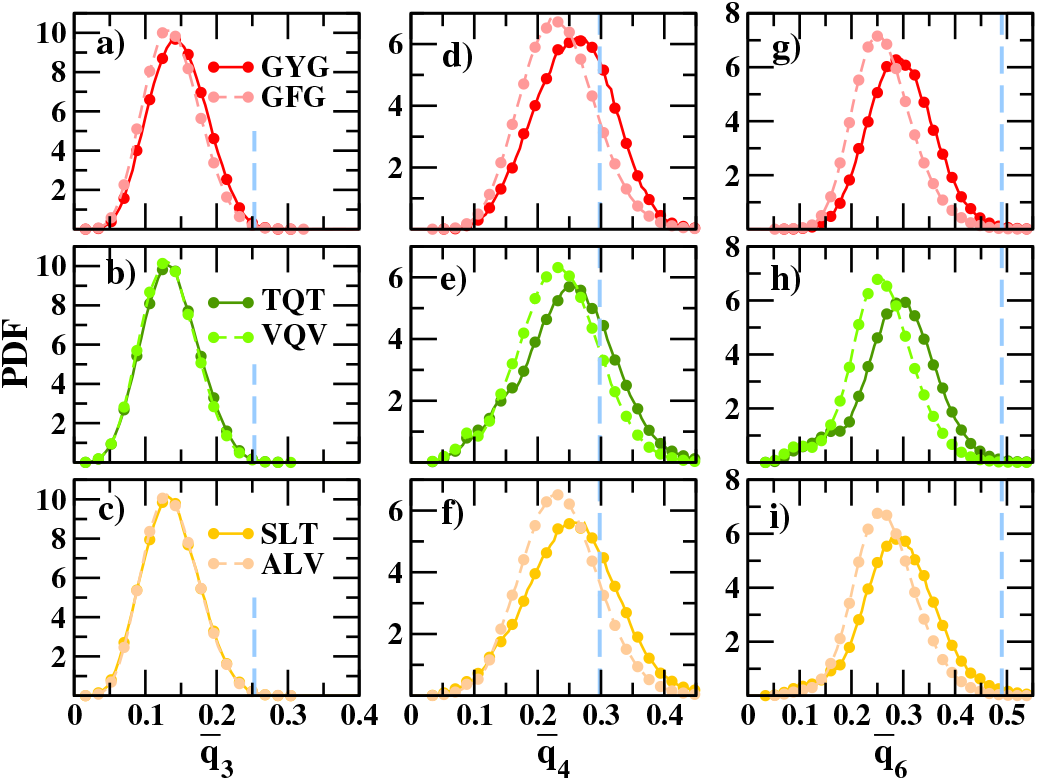
Probability density functions (PDF) of SBOPs for wild-type INP PINS and their mutated variant PINS. The top, middle, and bottom rows represent GYG-GFG, TQT-VQV, and SLT-ALV pairs, respectively. The three columns represents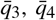, and 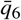 PDFs, respectively. The most probable location of the PDF for the hexagonal ice, I_*h*_, is shown using the dashed light blue line.

### B. GYG Enhances Tetrahedral Ordering of Interfacial Water toward an Ice-Like Tetrahedrality, Resembling Supercooled Water, More Strongly than Other PINS

The ensemble-averaged q_tet_ value, calculated using equation 5, equals 1 for a fully tetrahedral configuration and 0 for a completely disordered arrangement.^53–55^ Previous stud-ies have reported that hydrophilic amino acids disrupt the tetrahedral order of the hydration shell, whereas hydrophobic amino acids promote tetrahedrality.^53,56,57^ Among the wild-type PINS (Figure 4a), the probability density function (PDF) of q_tet_ for interfacial water (IW) near SGSE shows the broadest distribution, with a shoulder displaced from the *I*_*h*_ reference peak, indicating the least tetrahedrally ordered hydration shell among the four motifs consistent with the disruptive effect of the charged Glutamate (E) and polar Serine (S) residues.^58,59^ IWs at TQT show a modestly higher degree of tetrahedrality than SGSE. IWs at SLT show a peak position shifted toward lower q_tet_ relative to GYG, but a narrower, taller peak than both SGSE and TQT, indicating a larger fraction of IWs confined to a single, slightly distorted tetrahedral configuration rather than being broadly distributed across order-parameter space. The q_tet_ PDF for IWs at GYG shows a marked shift toward the *I*_*h*_ reference, together with the tallest and narrowest peak of the four motifs, indicating that GYG promotes tetrahedral order in its hydration shell more strongly than the other three motifs. The overall ranking of hydration-shell tetrahedrality is therefore GYG > SLT > TQT > SGSE.

**FIG. 4.**
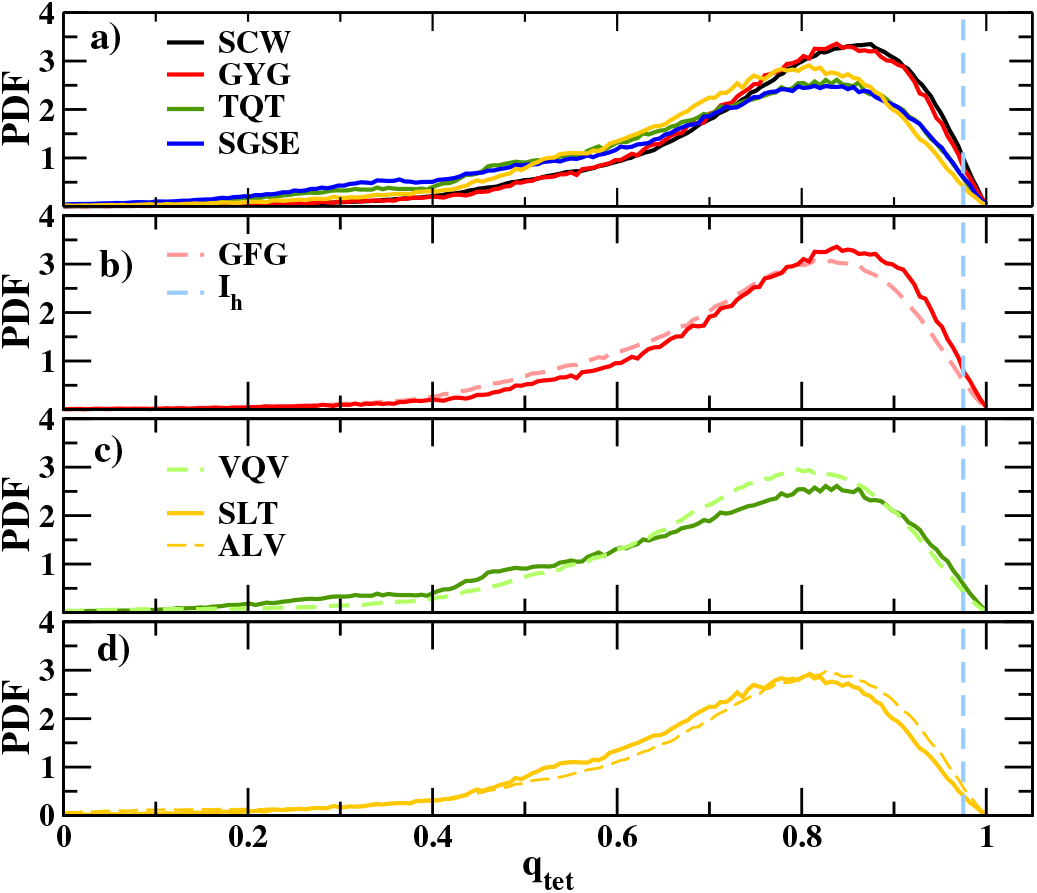
Probability density function (PDF) of the tetrahedral order parameter (q_*tet*_) of (a) all INP PINS, and the effect of mutation for (b) GYG-GFG, (c) TQT-VQV, and (d) SLT-ALV pair of PINS. The dashed light blue line shows the location of the peak of the probability density function of q_*tet*_ of bulk ice.

Next, the effect of ice-nucleating residues is examined using mutation. For GYG→GFG (Figure 4b), the GFG distribution showed a lower peak height and a shift toward lower q_tet_ relative to GYG, indicating reduced tetrahedral ordering of IWs at the mutant site. The reduction in tetrahedrality due to the Y to F mutation is attributed to the disruption in linearly arranged hydroxyl-mediated, ice-like structuring of IWs around the tyrosine residues rather than the hydrophobic effect.^53,56,57,59^ The TQT→VQV mutation results in a shift in the peak location of the probability distribution function (PDF) of *q*_*tet*_ away from that of the I_*h*_ with an increase in peak height. The SLT→ALV mutations result in increased IW tetrahedrality. The PDF of *q*_*tet*_ of ALV shows a shift towards the I_*h*_ PDF with a peak height comparable to SLT, indicating a shift towards higher tetrahedrality. This is due to the increasing side-chain hydrophobicity, which, in the absence of a hydroxyl group, enhances tetrahedral ordering of IWs as found earlier.^53,56,57,59^

### C. TQT Promotes Entropy of Interfacial Water Closest to Hexagonal Ice, Followed by GYG

The velocity autocorrelation function (VACF), density of states (DOS), and entropy of water molecules in bulk SCW, I_*h*_, and IWs at INP and mutated INP PINS are calculated using the 2PT method as mentioned in Method E of section III. 2PT is a very well-explored method to calculate various thermodynamic properties, especially entropy.^45,48,49,60^ The IW near all PINS have more pronounced negative minima in the translational VACF compared to SCW, and TQT has a slightly lower minimum compared to GYG (Figure 5a). The deeper minima in the translational VACF are indicative of enhanced backscattering, reflecting restricted local translational dynamics. Water molecules in the I_*h*_ structure, having the most constrained environment, show the deepest first minimum. The rotational VACF does not show any significant distinction between INP PINS and SCW, indicating that the rotational motion of water is not much hindered (Figure 5b). The difference between INP PINS, SCW, and I_*h*_ in total VACF is dominated by translational motion (Figure 5c). All the VACFs decay to zero by *t* ≈ 0.4 − 0.5 ps (data not shown). The translational DOS (5d) shows two peaks: the first peak at ≈ 50 cm^−1^ corresponds partly to H-bond bending and also signifies caging dynamics in dense fluids.^61^ The second peak at ≈ 250 cm^−1^ corresponds to H-bond stretching. The first peak of water near all INP PINS shows an ∼ 10 cm^−1^ blue shift and reduced height compared to SCW, indicating greater immobility of IW at PINS than in SCW. In rotational DOS (Figure 5e), a redshift indicates greater librational motion, and a blueshift indicates more constrained librational motion. The increased height means a less spread distribution, which corresponds to less orientational disorder. IWs at GYG and TQT PINS show the blue-shift, whereas IWs at SLT PINS show a slight red-shift compared to SCW. Also, the increase in peak height for IWs at PINS compared to SCW is the smallest for SLT, whereas GYG and TQT show a similar increase. Hence, IWs at SLT PINS exhibit greater librational freedom than those near GYG and TQT. Overall, GYG and TQT impose stronger translational and rotational constraints on IW than SLT. This is consistent with previous work emphasizing that the methyl group on threonine is essential for the structural rigidity required for ice nucleation, and mutating threonine to serine reduces motif rigidity, since the serine hydroxyl group rotates between two rotameric states and therefore cannot bind water effectively.^62^ The trends observed in the VACF and DOS distribution are directly reflected in the entropy values.

**FIG. 5.**
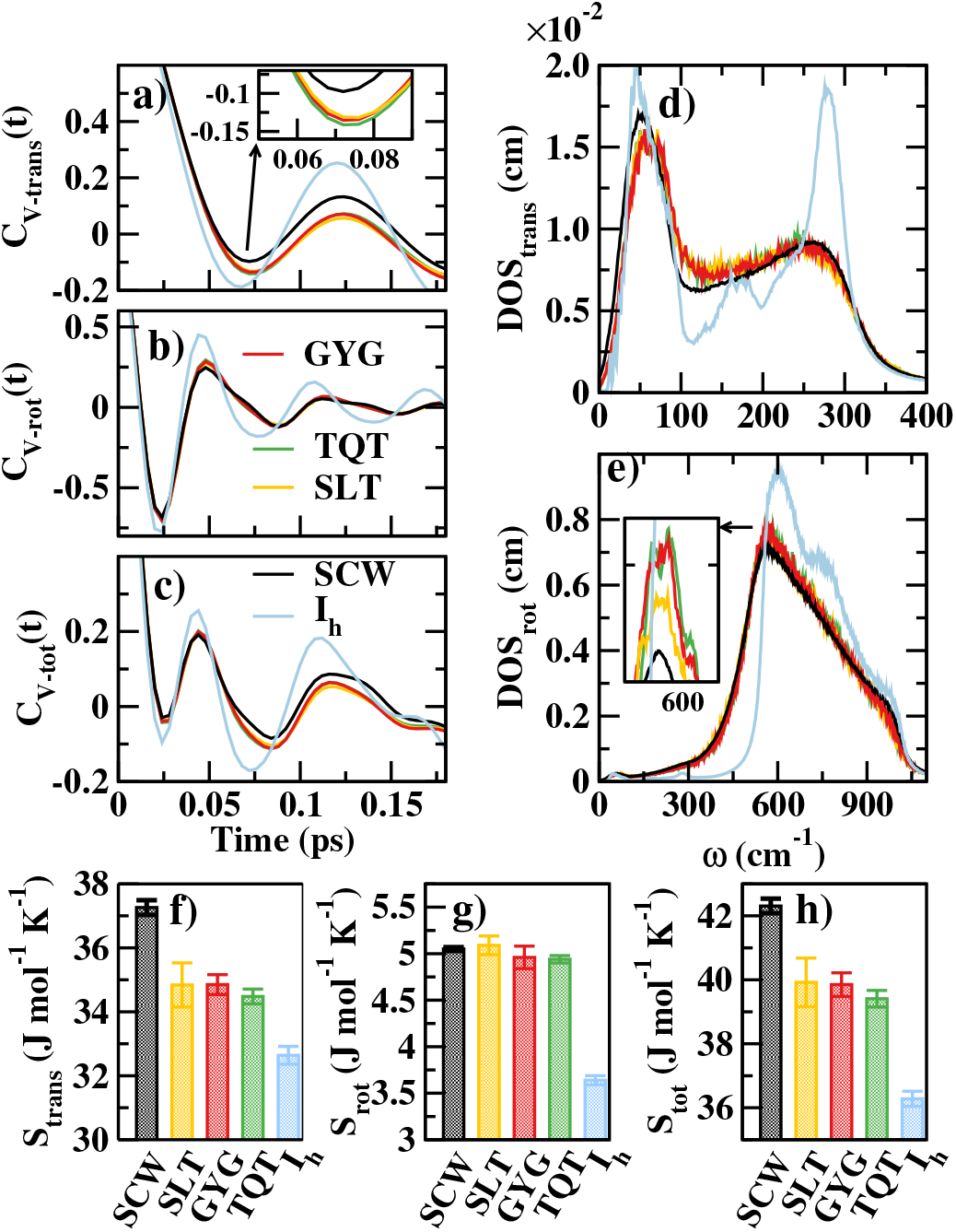
(a) Translational, (b) rotational, and (c) total VACF; (d) translational and (e) rotational density of states; (f) translational, (g) rotational, and (h) total entropy of SCW, IW near PINS, and I_*h*_.

The *S*_*trans*_, *S*_*rot*_, and *S*_*tot*_ of the IW near all PINS decrease with respect to the SCW. I_h_ has the least entropy among all (Figure 5f-h) due to the highly ordered water network of the hexagonal ice polymorph. The entropy difference, Δ*S* = *S*_SCW_ − *S*_PINS_ and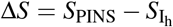, is shown in Tables S1 and S2 of the Supplementary Material. The reduction in entropy relative to SCW is largest for IWs near TQT, followed by GYG and SLT. The IW entropy is strongly modulated by the side-chain rigidity of TQT, due to the predominantly unimodal rotameric distribution of threonine residues, as seen in Figure S2a and S2c of the Supplementary Material. The rigidity imposes the largest constraints on the IW, resulting in the highest entropy reduction. GYG residues exhibit an intermediate entropy reduction due to fewer solvent-exposed water-anchoring motifs. In contrast, serine residues of SLT permit greater flexibility due to the bimodal rotameric distribution (Figure S2b and d of the Supplementary Material) and produce the least entropy reduction. These results highlight the critical role of side-chain rigidity in IW entropy reduction.^21,62,63^

The mutations of GYG and SLT do not have any significant change in the respective VACF of the associated IW (Figure 6a-b). Comparatively, IW near VQV shows a very minor difference after mutation (Figure 6c). Mutations in all PINS result in small differences in the density of states (Figure 6d-f) of the respective IW. A blueshift is observed for VQV and ALV relative to TQT and SLT, respectively, from 500 − 550cm^−1^ as shown in the inset. GFG shows a complete overlap in this region and a redshift beyond 550 cm^−1^. The blueshift is consistent with an increase in Δ*S* for mutations in SLT and TQT, where Δ*S* = *S*_*mut*_ − *S*_*wt*_, and *S*_*wt*_ and *S*_*mut*_ represent the entropy of IWs near the wild type and the mutated PINS, respectively. The translational, rotational, and total Δ*S* (Figure 6 g-i) increase from SLT→ALV and TQT→VQV, but decrease from GYG→GFG. The opposite behavior in Δ*S* depends on the interplay between hydrophobicity and local surface curvature. A qualitative structural explanation is that the increased hydrophobicity from GYG to GFG allows IW to pack and hydrogen-bond more tightly around the curved protrusive surface (Figure 1c) of the GYG PINS of the INP *β* helix, constraining dynamics and lowering entropy. In contrast, increased hydrophobicity in TQT and SLT due to mutation disrupts an already length-scale-limited hydration shell due to the flat surface occupied by TQT and SLT, with less slaved water dynamics and entropy. This trend is consistent with the length-scale framework proposed in earlier studies.^43^

**FIG. 6.**
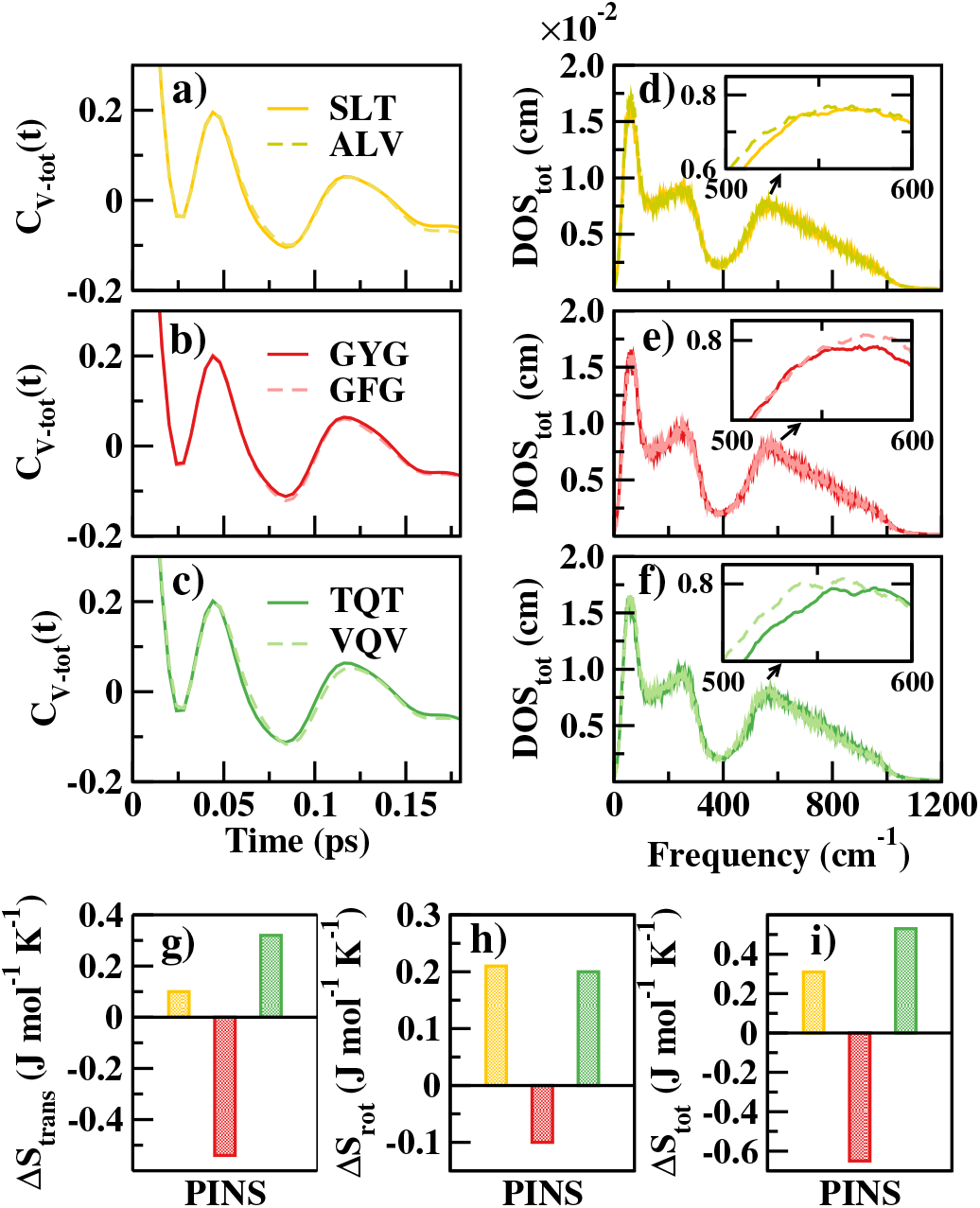
Comparison of total VACF for (a) SLT-ALV, (b) GYG-GFG, and (c) TQT-VQV pairs, and total density of states for (d) SLT-ALV, (e) GYG-GFG, (f) TQT-VQV. Change in (g) translational, (h) rotational, and (i) total entropy upon mutation of INP PINS.

## IV. CONCLUSIONS

Bacterial ice nucleating proteins (INPs) are the most active biological heterogeneous ice nucleators, with direct relevance to global climate due to their extreme environmental resilience and abundance. Molecular-level elucidation of the specific ice-nucleation sites on INPs and the structural and interfacial determinants governing their activity through molecular simulations is crucial for establishing the mechanistic basis of biological ice nucleation and enabling the rational design of efficient, tunable ice-nucleating materials. Previous studies^15,16,20^ have predicted the highly conserved TXT and SXT motifs in INPs as potential ice-nucleation sites, based on their proposed ability to organize IW into ice-like templates on relatively flat protein surfaces. However, a critical gap remains, as these studies have not systematically explored whether these motifs represent the most potent nucleation sites among the most conserved sequence motifs on the four surfaces of the model INP. Specific sequence features and corresponding interfacial structural environments that are most conducive to ice nucleation are not well established. The functions of motifs other than TXT remain largely overlooked. This limits the rational exploitation in future protein engineering. To this end, all-atom molecular dynamics simulations of the SCW, bulk I_*h*_, INP monomer, its mutants are performed using the CHARMM36m force field and the TIP4P/Ice water model at 260 K, below the ice nucleation temperature of the INP monomer. Using the well-established sequence^20^, a 16-loop *β* -helical protein is modeled using Alphfold-3. Four potential ice-nucleating sites (PINS) on four surfaces have been identified: GYG, TQT, SGSE, and SLT. To verify the role of the defined PINS in ice nucleation, the key residues of the PINS are mutated by either removing hydroxyl groups or replacing them with methyl groups, converting the GYG, TQT, and SLT PINS into hydrophobic GFG, VQV, and ALV PINS. Randomly oriented tyrosine residues from the GYG sites of the wild-type INP monomer align during simulation to form a phenyl-ring-stacked, ladder-like structure, where hydroxyl groups are present in a linear fashion. The PDFs of 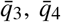, and 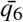 SBOPs of the wild-type INPs demonstrate that the probability for the IW associated with GYG is closest to that of the *I*_*h*_, compared to TQT or SLT, and the probability is farthest for SGSE. All the mutated PINS show a lower order compared to the wild-type PINS, with the effect most prominent for the GYG-GFG pair. The effect of mutation is more significant for 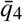 and 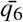 compared to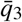, suggesting that 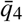and 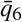 are better parameters to monitor the effect of residue mutation on the local structural bond order of IW molecules. The probability of tetrahedrality of the IW increases from SGSE to TQT or SLT to GYG, similar to the trend found in the SBOP. GYG imposing the highest structural order in the IW compared to conventional TQT or SLT motif is in similar line to the previous experimental prediction that the threonine-based motif does not induce greater ordering of water than other PINS.^52^

Although the hydrophilic-to-hydrophobic mutations increase the tetrahedrality of IWs at ALV PINS, the GYG-to-GFG mutation shows the opposite behavior. GYG mutation reduces IW tetrahedrality by disrupting hydroxyl-mediated ice-like ordering, whereas TQT and SLT increase tetrahedrality through enhanced side-chain hydrophobicity in the absence of hydroxyl groups, similar to a previous report.^53,56,57,59^ 2PT analysis shows that the IW entropy at all INP PINS is reduced relative to SCW, reflecting the IW confinement. TQT exhibits the lowest IW entropy, closest to ice I_*h*_, followed by GYG, whereas SLT shows the highest entropy due to greater serine rotameric flexibility. Mutational analyses further demonstrate that IW entropy is governed by the coupled effects of surface geometry, hydrophobicity, and water-anchoring motifs: GYG to GFG mutation enhances water packing on the curved hydrophobic surface, whereas increased hydrophobicity on flat TQT and SLT surfaces disrupts local hydration structure. Overall, Y is the primary residue in the GYG motif that has the maximum water-ordering capacity among all residues of all PINS.

Our results indicate that entropic reduction alone is insufficient to explain the emergence and stabilization of ice-like ordering within the IW layer. Instead, Steinhardt bond ordering in the IW proximal to the PINS provides a critical measure of the most potent ice nucleator. The ice-like water ordering is collectively governed by geometric compatibility between ice-nucleating sites and the ice-like lattice, along with a reduction in IW entropy toward ice-like values, side-chain hydrophobicity, interfacial geometry, and previously explored sequence conservation.^13,15,20^ By systematically characterizing the interfacial ice-nucleating surface of the INP into four PINS, our results reveal that GYG as an unrecognized and probably a better contributor to ice-like hydration ordering, in conjunction with the previously proposed TQT and SLT motifs.^20^ Our conclusion converges with a parallel, independently conducted, very recent nucleation-kinetics study, which finds that ice growth in a model INP initiates predominantly from the YGS motif and that increasing YGS/TXT density enhances nucleation efficiency.^64^ That work, however, characterizes the contribution of motifs through ice-nucleation propagation and hydrogen-bonding kinetics in a spontaneously freezing system, leaving open the structural and thermodynamic basis for why GYG-associated water is more ice-like than water near the canonical threonine motif^64^. Our work provides mechanistic details and the thermodynamic basis of the ice-nucleation ability of the GYG motif in INP and quantifies the structural order around different sites in INP. Although the enhanced order alone is insufficient for ice nucleation and growth at 260 *K*, underscoring the requirement for INP dimerization in future investigation, the current study proposes a revised molecular framework that provides the foundation for protein-mediated ice nucleation. Beyond resolving the key mechanistic question of biological ice nucleation, these insights identify structural features that can be rationally targeted to enhance nucleation efficiency, thereby enabling the engineering of next-generation ice-nucleating proteins with improved performance for applications in biotechnology, cryopreservation, and climate-related technologies.

## Supporting information

See the supplementary material for a plot of the radial dis- tribution function, tables of entropy difference between bulk phases and interface water,

## SUPPLEMENTARY MATERIAL

See the supplementary material for a plot of the radial distribution function, tables of entropy difference between bulk phases and interface water, and rotameric states of threonine and serine residues.

## ACKNOWLEDGMENTS

This research is supported by the Department of Biotechnology Bio-RIDE scheme (Grant ID: BT/PR65037/MRNAR/169/43/2026), Ministry of Science and Technology, Government of India.

## AUTHOR DECLARATIONS

### Conflict of Interest

The authors have no conflicts to disclose.

## Author Contributions

**Jadhav, S. E**.: Validation, Formal analysis, Investigation, Data curation, Writing — original draft, Visualization. **Debnath, A**.: Conceptualization, Validation, Formal analysis, Resources, Writing — review & editing, Supervision, Project administration, Funding acquisition.

## DATA AVAILABILITY STATEMENT

The data that support the findings of this study are available within the article [and its supplementary material]. Additional data are openly available in the GitHub repository https://github.com/sankalpgit1/INP_mutations_par.

## Notes

### Competing Interest Statement

The authors have declared no competing interest.

## References

1 J. D. Atkinson, B. J. Murray, M. T. Woodhouse, T. F. Whale, K. J. Baustian, K. S. Carslaw, S. Dobbie, D. O’Sullivan, and T. L. Malkin, “The importance of feldspar for ice nucleation by mineral dust in mixed-phase clouds,” Nature 498, 355–358 (2013).

2 T. Kawasaki and H. Tanaka, “Formation of a crystal nucleus from liquid,” Proceedings of the National Academy of Sciences 107, 14036–14041 (2010).

3 P. J. Ward and P. J. DeMott, “Preliminary experimental evaluation of sno-max (tm) snow inducer, nucleus pseudomonas syringae, as an artificial ice for weather modification,” The Journal of Weather Modification 21, 1 (1989).

4 B. Murray, S. Broadley, T. Wilson, S. Bull, R. Wills, H. Christenson, and E. Murray, “Kinetics of the homogeneous freezing of water,” Physical Chemistry Chemical Physics 12, 10380–10387 (2010).

5 J. Bühl, A. Ansmann, P. Seifert, H. Baars, and R. Engelmann, “Toward a quantitative characterization of heterogeneous ice formation with lidar/radar: Comparison of calipso/cloudsat with ground-based observations,” Geophysical Research Letters 40, 4404–4408 (2013).

6 H. Laksmono, T. A. McQueen, J. A. Sellberg, N. D. Loh, C. Huang, D. Schlesinger, R. G. Sierra, C. Y. Hampton, D. Nordlund, M. Beye, A. V. Martin, A. Barty, M. M. Seibert, M. Messerschmidt, G. J. Williams, S. Boutet, K. Amann-Winkel, T. Loerting, L. G. M. Pettersson, M. J. Bogan, and A. Nilsson, “Anomalous behavior of the homogeneous ice nucleation rate in “no-man’s land”,” The Journal of Physical Chemistry Letters 6, 2826–2832 (2015).

7 D. Gurian-Sherman and S. E. Lindow, “Bacterial ice nucleation: significance and molecular basis,” The FASEB Journal 7, 1338–1343 (1993).

8 R. L. Green and G. J. Warren, “Physical and functional repetition in a bacterial ice nucleation gene,” Nature 317, 645–648 (1985).

9 S. E. Lindow, D. C. Arny, and C. D. Upper, “Bacterial ice nucleation: a factor in frost injury to plants,” Plant Physiology 70, 1084–1089 (1982).

10 M. Lukas, R. Schwidetzky, R. J. Eufemio, M. Bonn, and K. Meister, “Toward understanding bacterial ice nucleation,” The Journal of Physical Chemistry B 126, 1861–1867 (2022).

11 P. Amato, M. Joly, C. Schaupp, E. Attard, O. Möhler, C. E. Morris, Y. Brunet, and A.-M. Delort, “Survival and ice nucleation activity of bacteria as aerosols in a cloud simulation chamber,” Atmospheric Chemistry and Physics 15, 6455–6465 (2015).

12 C. E. Morris, F. Conen, J. Alex Huffman, V. Phillips, U. Pöschl, and D. C. Sands, “Bioprecipitation: a feedback cycle linking earth history, ecosystem dynamics and land use through biological ice nucleators in the atmosphere,” Global Change Biology 20, 341–351 (2014).

13 Y. Qiu, A. Hudait, and V. Molinero, “How size and aggregation of ice-binding proteins control their ice nucleation efficiency,” Journal of the American Chemical Society 141, 7439–7452 (2019).

14 R. Schwidetzky, P. Sudera, A. T. Backes, U. Poschl, M. Bonn, J. Frohlich-Nowoisky, and K. Meister, “Membranes are decisive for maximum freezing efficiency of bacterial ice nucleators,” The Journal of Physical Chemistry Letters 12, 10783–10787 (2021).

15 C. P. Garnham, R. L. Campbell, V. K. Walker, and P. L. Davies, “Novel dimeric β-helical model of an ice nucleation protein with bridged active sites,” BMC Structural Biology 11, 36 (2011).

16 R. Aich, P. Pal, S. Chakraborty, and B. Jana, “Preferential ordering and organization of hydration water favor nucleation of ice by ice-nucleating proteins over antifreeze proteins,” The Journal of Physical Chemistry B 127, 6038–6048 (2023).

17 A. K. Shargh, C. D. Stiles, and J. A. El-Awady, “Effect of ice nucleating proteins on the structure-property relationships of ice: A molecular dynamics study,” The Journal of Chemical Physics 164, 024106 (2026).

18 S. Hartmann, M. Ling, L. S. A. Dreyer, A. Zipori, K. Finster, S. Grawe, L. Z. Jensen, S. Borck, N. Reicher, T. Drace, D. Niedermeier, N. C. Jones, S. V. Hoffmann, H. Wex, Y. Rudich, T. Boesen, and T. Šantl Temkiv, “Structure and protein-protein interactions of ice nucleation proteins drive their activity,” Frontiers in Microbiology 13, 872306 (2022).

19 S. J. Roeters, T. W. Golbek, M. Bregnhøj, T. Drace, S. Alamdari, W. Roseboom, G. Kramer, T. Šantl-Temkiv, K. Finster, J. Pfaendtner, S. Woutersen, T. Boesen, and T. Weidner, “Ice-nucleating proteins are activated by low temperatures to control the structure of interfacial water,” Nature Communications 12, 1183 (2021).

20 A. Hudait, N. Odendahl, Y. Qiu, F. Paesani, and V. Molinero, “Ice-nucleating and antifreeze proteins recognize ice through a diversity of anchored clathrate and ice-like motifs,” Journal of the American Chemical Society 140, 4905–4912 (2018).

21 D. Saha, R. Aich, A. Mukherjee, and B. Jana, “Water entropy at the threonine-rich surface of antifreeze and ice-nucleating proteins: small changes make a big difference,” Chemical Science 16, 10771–10784 (2025).

22 J. Forbes, A. Bissoyi, L. Eickhoff, N. Reicher, T. Hansen, C. G. Bon, K. Walker, T. Koop, Y. Rudich, I. Braslavsky, and P. L. Davies, “Water-organizing motif continuity is critical for potent ice nucleation protein activity,” Nature Communications 13, 5019 (2022).

23 M. Lukas, R. Schwidetzky, A. T. Kunert, E. H. Backus, U. Poschl, J. Frohlich-Nowoisky, M. Bonn, and K. Meister, “Interfacial water ordering is insufficient to explain ice-nucleating protein activity,” The Journal of Physical Chemistry Letters 12, 218–223 (2021).

24 P. Bieber and N. Borduas-Dedekind, “High-speed cryo-microscopy reveals that ice-nucleating proteins of pseudomonas syringae trigger freezing at hydrophobic interfaces,” Science Advances 10, eadn6606 (2024).

25 K. Basu, L. A. Graham, R. L. Campbell, and P. L. Davies, “Flies expand the repertoire of protein structures that bind ice,” Proceedings of the National Academy of Sciences 112, 737–742 (2015).

26 K. Basu, S. S. Wasserman, P. S. Jeronimo, L. A. Graham, and P. L. Davies, “Intermediate activity of midge antifreeze protein is due to a tyrosine-rich ice-binding site and atypical ice plane affinity,” The FEBS Journal 283, 1504–1515 (2016).

27 J. Abramson, J. Adler, J. Dunger, R. Evans, T. Green, A. Pritzel, O. Ronneberger, L. Willmore, A. J. Ballard, J. Bambrick, S. W. Bodenstein, D. A. Evans, C.-C. Hung, M. O’Neill, D. Reiman, K. Tunyasuvunakool, Z. Wu, A. Žemgulytė, E. Arvaniti, C. Beattie, O. Bertolli, A. Bridgland, A. Cherepanov, M. Congreve, A. I. Cowen-Rivers, A. Cowie, M. Figurnov, F. B. Fuchs, H. Gladman, R. Jain, Y. A. Khan, C. M. R. Low, K. Perlin, A. Potapenko, P. Savy, S. Singh, A. Stecula, A. Thillaisundaram, C. Tong, S. Yakneen, E. D. Zhong, M. Zielinski, A. Žídek, V. Bapst, P. Kohli, M. Jaderberg, D. Hassabis, and J. M. Jumper, “Accurate structure prediction of biomolecular interactions with alphafold 3,” Nature 630, 493–500 (2024).

28 M. Ling, H. Wex, S. Grawe, J. Jakobsson, J. Löndahl, S. Hartmann, K. Finster, T. Boesen, and T. Šantl-Temkiv, “Effects of ice nucleation protein repeat number and oligomerization level on ice nucleation activity,” Journal of Geophysical Research: Atmospheres 123, 1802–1810 (2018).

29 J. Lee, X. Cheng, S. Jo, A. D. MacKerell, J. B. Klauda, and W. Im, “Charmm-gui input generator for namd, gromacs, amber, openmm, and charmm/openmm simulations using the charmm36 additive force field,” Biophysical journal 110, 641a (2016).

30 B. R. Brooks, C. L. Brooks III, A. D. Mackerell Jr., L. Nilsson, R. J. Petrella, B. Roux, Y. Won, G. Archontis, C. Bartels, S. Boresch, A. Caflisch, L. Caves, Q. Cui, A. R. Dinner, M. Feig, S. Fischer, J. Gao, M. Hodoscek, W. Im, K. Kuczera, T. Lazaridis, J. Ma, V. Ovchinnikov, E. Paci, R. W. Pastor, C. B. Post, J. Z. Pu, M. Schaefer, B. Tidor, R. M. Venable, H. L. Woodcock, X. Wu, W. Yang, D. M. York, and M. Karplus, “Charmm: the biomolecular simulation program,” Journal of Computational Chemistry 30, 1545–1614 (2009).

31 S. Jo, T. Kim, V. G. Iyer, and W. Im, “Charmm-gui: a web-based graphical user interface for charmm,” Journal of Computational Chemistry 29, 1859–1865 (2008).

32 J. Abascal, E. Sanz, R. García Fernández, and C. Vega, “A potential model for the study of ices and amorphous water: Tip4p/ice,” The Journal of Chemical Physics 122, 234511 (2005).

33 J. Huang, S. Rauscher, G. Nawrocki, T. Ran, M. Feig, B. L. De Groot, H. Grubmüller, and A. D. MacKerell Jr, “Charmm36m: an improved force field for folded and intrinsically disordered proteins,” Nature Methods 14, 71–73 (2017).

34 Y. Wardi, “A stochastic steepest-descent algorithm,” Journal of Optimization Theory and Applications 59, 307–323 (1988).

35 H. J. Berendsen, J. v. Postma, W. F. Van Gunsteren, A. DiNola, and J. R. Haak, “Molecular dynamics with coupling to an external bath,” The Journal of Chemical Physics 81, 3684–3690 (1984).

36 M. Parrinello and A. Rahman, “Polymorphic transitions in single crystals: A new molecular dynamics method,” Journal of Applied Physics 52, 7182–7190 (1981).

37 G. Bussi, D. Donadio, and M. Parrinello, “Canonical sampling through velocity rescaling,” The Journal of Chemical Physics 126, 014101 (2007).

38 U. Essmann, L. Perera, M. L. Berkowitz, T. Darden, H. Lee, and L. G. Pedersen, “A smooth particle mesh ewald method,” The Journal of Chemical Physics 103, 8577–8593 (1995).

39 M. Matsumoto, T. Yagasaki, and H. Tanaka, “Genice: hydrogen-disordered ice generator,” Journal of Computational Chemistry 39, 61–64 (2018).

40 P. J. Steinhardt, D. R. Nelson, and M. Ronchetti, “Bond-orientational order in liquids and glasses,” Physical Review B 28, 784–805 (1983).

41 W. Lechner and C. Dellago, “Accurate determination of crystal structures based on averaged local bond order parameters,” The Journal of Chemical Physics 129, 114707 (2008).

42 J. R. Errington and P. G. Debenedetti, “Relationship between structural order and the anomalies of liquid water,” Nature 409, 218–321 (2001).

43 V. R. Hande and S. Chakrabarty, “Structural order of water molecules around hydrophobic solutes: length-scale dependence and solute–solvent coupling,” The Journal of Physical Chemistry B 119, 11346–11357 (2015).

44 S.-T. Lin, M. Blanco, and W. A. Goddard III, “The two-phase model for calculating thermodynamic properties of liquids from molecular dynamics: Validation for the phase diagram of lennard-jones fluids,” The Journal of Chemical Physics 119, 11792–11805 (2003).

45 S.-T. Lin, P. K. Maiti, and W. A. Goddard III, “Two-phase thermodynamic model for efficient and accurate absolute entropy of water from molecular dynamics simulations,” The Journal of Physical Chemistry B 114, 8191–8198 (2010).

46 T. A. Pascal, W. A. Goddard, and Y. Jung, “Entropy and the driving force for the filling of carbon nanotubes with water,” Proceedings of the National Academy of Sciences 108, 11794–11798 (2011).

47 C.-H. Ho and F. Paesani, “Entropy of liquid water as predicted by the two-phase thermodynamic model and data-driven many-body potentials,” The Journal of Physical Chemistry B 128, 6885–6891 (2024).

48 A. Debnath, K. Ayappa, and P. K. Maiti, “Simulation of influence of bilayer melting on dynamics and thermodynamics of interfacial water,” Physical Review Letters 110, 018303 (2013).

49 A. Debnath, B. Mukherjee, K. Ayappa, P. K. Maiti, and S.-T. Lin, “Entropy and dynamics of water in hydration layers of a bilayer,” The Journal of Chemical Physics 133, 174704 (2010).

50 K. Olechnovič and S. Grudinin, “Voronota-lt: Efficient, flexible, and solvent-aware tessellation-based analysis of atomic interactions,” Journal of Computational Chemistry 46, e70178 (2025).

51 J. Shi, M. Fulford, H. Li, M. Marzook, M. Reisjalali, M. Salvalaglio, and C. Molteni, “Investigating the quasi-liquid layer on ice surfaces: a comparison of order parameters,” Physical Chemistry Chemical Physics 24, 12476–12487 (2022).

52 R. Pandey, K. Usui, R. A. Livingstone, S. A. Fischer, J. Pfaendtner, E. H. G. Backus, Y. Nagata, J. Fröhlich-Nowoisky, L. Schmüser, S. Mauri, J. F. Scheel, D. A. Knopf, U. Pöschl, M. Bonn, and T. Weidner, “Ice-nucleating bacteria control the order and dynamics of interfacial water,” Science Advances 2, e1501630 (2016).

53 D. R. Nutt and J. C. Smith, “Dual function of the hydration layer around an antifreeze protein revealed by atomistic molecular dynamics simulations,” Journal of the American Chemical Society 130, 13066–13073 (2008).

54 I. Gladich and M. Roeselová, “Comparison of selected polarizable and non-polarizable water models in molecular dynamics simulations of ice i h,” Physical Chemistry Chemical Physics 14, 11371–11385 (2012).

55 M. Riera, E. Lambros, T. T. Nguyen, A. W. Götz, and F. Paesani, “Low-order many-body interactions determine the local structure of liquid water,” Chemical Science 10, 8211–8218 (2019).

56 Z. F. Brotzakis, I. K. Voets, H. J. Bakker, and P. G. Bolhuis, “Water structure and dynamics in the hydration layer of a type iii anti-freeze protein,” Physical Chemistry Chemical Physics 20, 6996–7006 (2018).

57 S. A. Deshmukh, L. A. Solomon, G. Kamath, H. C. Fry, and S. K. Sankaranarayanan, “Water ordering controls the dynamic equilibrium of micelle– fibre formation in self-assembly of peptide amphiphiles,” Nature Communications 7, 12367 (2016).

58 M. Ide, Y. Maeda, and H. Kitano, “Effect of hydrophobicity of amino acids on the structure of water,” The Journal of Physical Chemistry B 101, 7022–7026 (1997).

59 K. Shiraga, T. Suzuki, N. Kondo, and Y. Ogawa, “Hydration and hydrogen bond network of water around hydrophobic surface investigated by terahertz spectroscopy,” The Journal of Chemical Physics 141, 235103 (2014).

60 T. A. Pascal, S.-T. Lin, and W. A. Goddard III, “Thermodynamics of liquids: standard molar entropies and heat capacities of common solvents from 2pt molecular dynamics,” Physical Chemistry Chemical Physics 13, 169–181 (2010).

61 J. Martínez Cordeiro and N. Aluru, “Water at low frequencies: Vibrational decomposition reveals spectral fingerprints of surface charge and dynamic surfaces,” The Journal of Chemical Physics 164, 164305 (2026).

62 A. Hudait, Y. Qiu, N. Odendahl, and V. Molinero, “Hydrogen-bonding and hydrophobic groups contribute equally to the binding of hyperactive antifreeze and ice-nucleating proteins to ice,” Journal of the American Chemical Society 141, 7887–7898 (2019).

63 U. S. Midya and S. Bandyopadhyay, “Elucidating the sluggish water dynamics at the ice-binding surface of the hyperactive tenebrio molitor antifreeze protein,” The Journal of Physical Chemistry B 127, 121–132 (2023).

64 E. A. Cino and D. P. Tieleman, “Simulation guided design of a potentially hyperactive ice nucleating protein,” Journal of Chemical Information and Modeling (2026).

