## Supplementary material for "Nanoscopic Origin of Enhanced Tyrosine-Mediated Water Ordering over Canonical Threonine Motifs in a Bacterial Ice-Nucleating Protein": See the supplementary material for a plot of the radial dis- tribution function, tables of entropy difference between bulk phases and interface water,

### 1 Radial Distribution Function

The radial distribution function (RDF) gives the probability of finding a particle at a given location and is calculated as

$$g(r) = \frac{1}{\rho_N 4\pi r^2} \langle \sum_{\substack{i,j \\ i \neq j}}^{N} \delta(r_{ij} - r) \rangle. \quad (1)$$

where $\rho_N$ is the number density of the bulk, r denotes the radius of the sphere, and $r_{ij}$ is the distance between $i^{th}$ and $j^{th}$ particles in the simulation box. The RDF is calculated between water oxygen atoms and all the heavy sidechain atoms (excluding hydrogen and $C_\alpha$ atoms) of the INP and mutated INP PINS. Figure S1(a) shows the RDF for water with reference to INP PINS. It is seen that the height of the first shell is proportional to the number of water-facing polar groups in the side chain. For GYG PINS, there's only one polar OH motif, so it shows the lowest first peak height. For TQT and SLT, both have two OH motifs that face the water and thus have the same first peak height, which is larger than that of GYG. SGSE has the highest peak because it has three water-facing polar groups, and also a charge that can hold water more strongly. Due to the charge, the RDF of SGSE is slightly left-shifted compared to other PINS. Among all the PINS, water from the GYG surface exhibits long-range order up to four shells. For other motifs, the weak long-range order is visible up to three hydration shells. For all PINS, the first shell is approximately 0.32 nm. The Figure S1 (b-d) shows the comparison of RDF for water from GYG vs GFG, TQT vs VQV, and SLT vs ALV references, respectively. Because mutated sites lack polar groups, their first hydration shell minimum is around 0.45, corresponding to the second hydration shell for INP PINS. Mutated INP PINS do not show any long-range hydration shell structure due to the lack of polar groups

### 2 Entropy Difference Between Bulk Phases and Interface Water

Table S1: Entropy difference $\Delta S = S_{\mathrm{SCW}} - S_{\mathrm{PINS}}$ for IWs near the GYG, TQT, and SLT PINS

| **PINS** | $\Delta S_{\mathrm{trans}}$ | $\Delta S_{\mathrm{rot}}$ | $\Delta S_{\mathrm{tot}}$ |
|---|---|---|---|
| TQT | 2.78 | 0.11 | 2.91 |
| GYG | 2.41 | 0.05 | 2.47 |
| SLT | 2.42 | -0.04 | 2.40 |

Table S2: Entropy difference $\Delta S = S_{\mathrm{PINS}} - S_{\mathrm{I_h}}$ for IWs near the GYG, TQT, and SLT PINS

| **PINS** | $\Delta S_{\mathrm{trans}}$ | $\Delta S_{\mathrm{rot}}$ | $\Delta S_{\mathrm{tot}}$ |
|---|---|---|---|
| TQT | 1.84 | 1.30 | 3.13 |
| GYG | 2.21 | 1.36 | 3.57 |
| SLT | 2.20 | 1.45 | 3.64 |

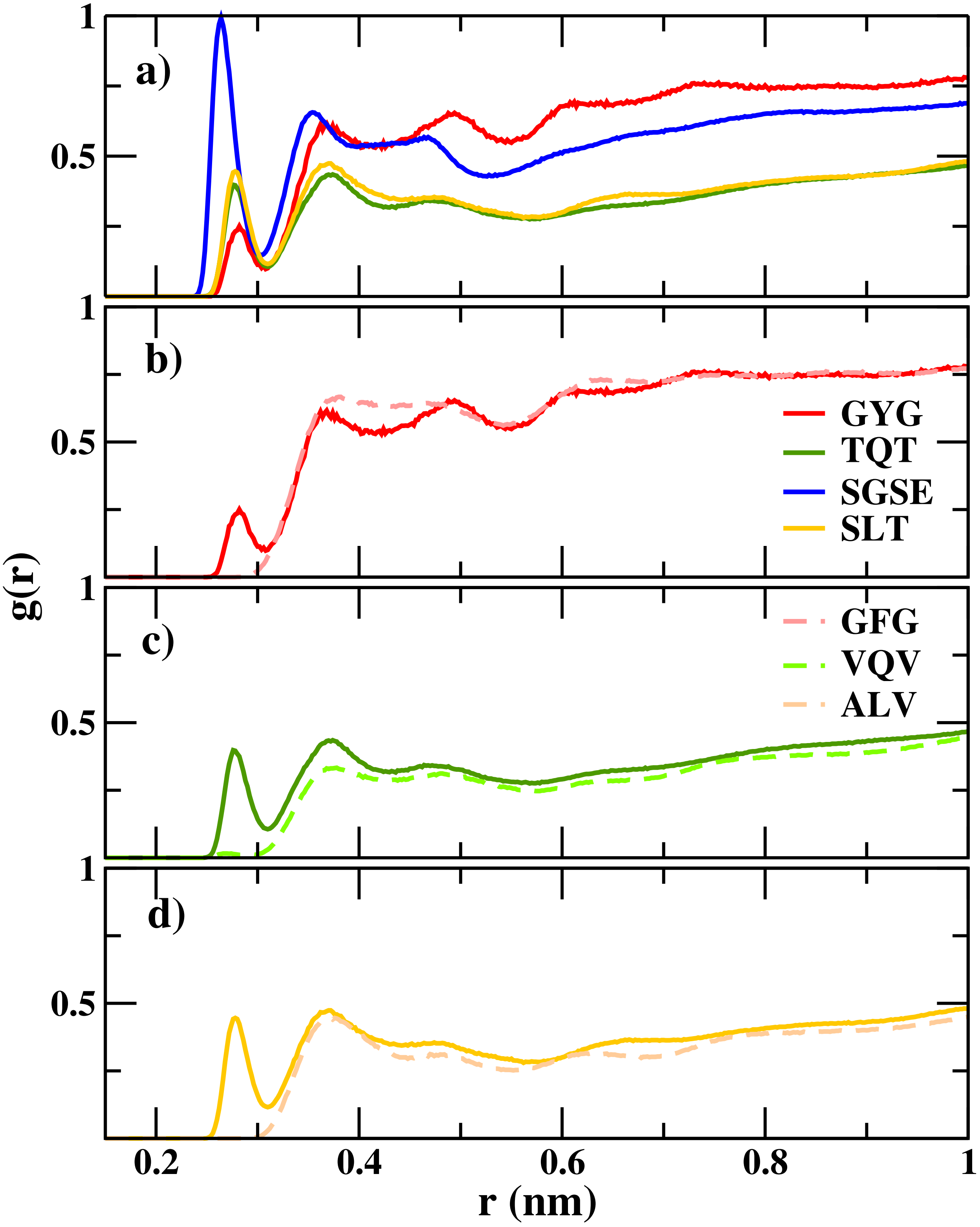


Figure S1: Radial distribution function (RDF) for a) all PINS of INP; comparison of (b) GYG vs GFG, (c) TQT vs VQV, (d) SLT vs ALV.

#### 3 Rotameric States for Serine and Threonine Residues

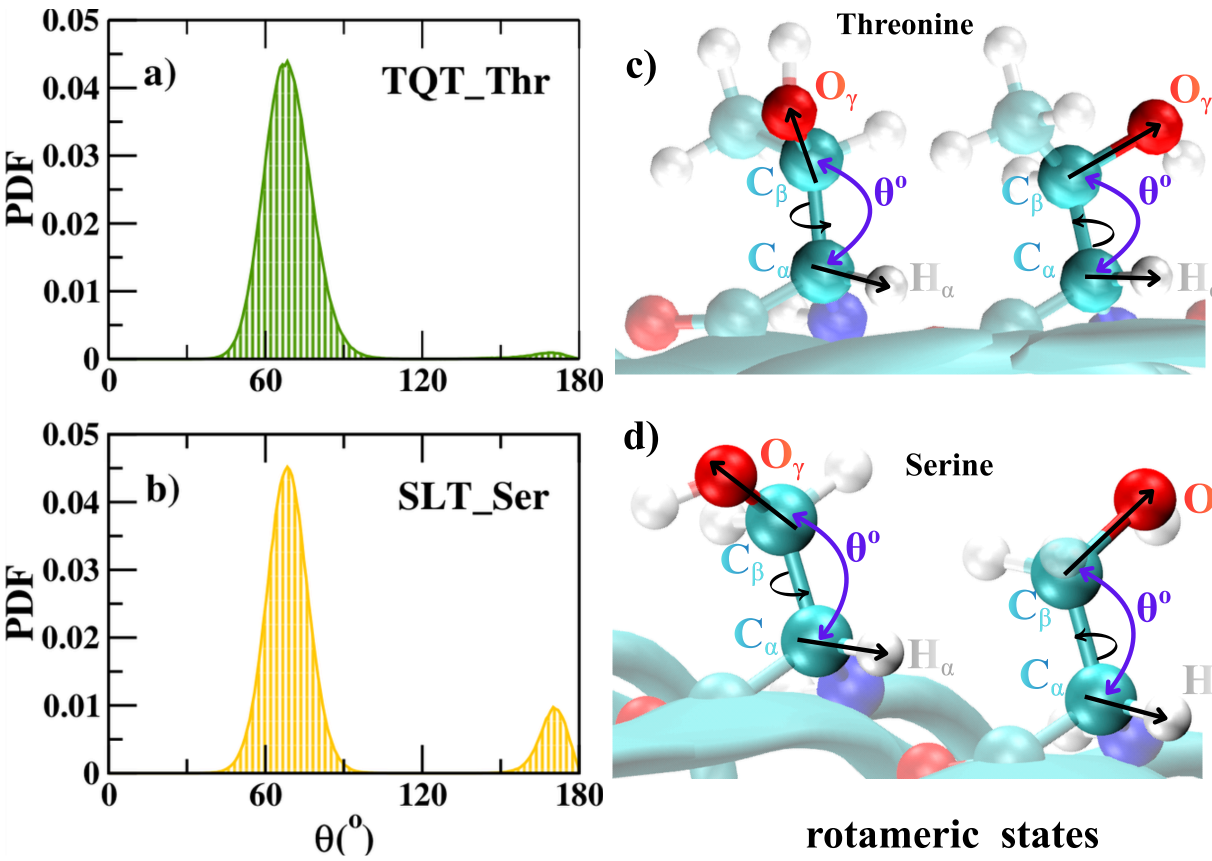


Figure S2: Probability distribution function of the angle between the $C_\alpha$-$H_\alpha$ and $C_\beta$-$O_\gamma$ vectors of (a) one of the threonine residues of TQT and (b) the serine residue of SLT. The pictorial representation of vectors for two rotameric states is shown for (c) threonine and (d) serine.
